# Accurate typing of class I human leukocyte antigen by Oxford nanopore sequencing

**DOI:** 10.1101/178590

**Authors:** Chang Liu, Fangzhou Xiao, Jessica Hoisington-Lopez, Kathrin Lang, Philipp Quenzel, Brian Duffy, Robi David Mitra

## Abstract

Oxford Nanopore Technologies’ MinION has expanded the current DNA sequencing toolkit by delivering long read lengths and extreme portability. The MinION has the potential to enable expedited point-of-care human leukocyte antigen (HLA) typing, an assay routinely used to assess the immunological compatibility between organ donors and recipients, but the platform’s high error rate makes it challenging to type alleles with clinical-grade accuracy. Here, we developed and validated Athlon, an algorithm that iteratively scores nanopore reads mapped to a hierarchical database of HLA alleles to arrive at a consensus diploid genotype; Athlon achieved a 100% accuracy in class I HLA typing at high resolution.

## MAIN TEXT

The Oxford Nanopore Technologies’ (ONT) MinION is a portable device the size of a mobile phone that performs rapid single-molecule sequencing.^1^ This device directly records dynamic changes in electric current across a nanopore as a single-stranded DNA or RNA molecule is ratcheted through the pore by a motor protein. The raw signals from hundreds of working nanopores are converted to sequencing reads in real time via an online base caller. The portability of this system, combined with its superior read lengths of up to 50 kb^2^, make the MinION uniquely positioned to enable point-of-care clinical sequencing, especially for applications that require haplotype information.

Advances in flow cell design and base-calling algorithms have led to steady improvements in the MinION’s raw read accuracy, which was as low as 66% in early 2014^3^ and is now approximately 92%^4^. However, despite initial successes in diagnostic microbiology,^5-10^ the relatively high error rate and the lack of a dedicated variant caller for diploid genomes have prevented the MinION from achieving widespread use in human DNA sequencing.^3^, ^11^ Here, we report a method for the targeted nanopore sequencing of class I HLA genes and a bioinformatic pipeline to interpret these data (Athlon; <u>http://github.com/cliu32/Athlon</u>). Together, these provide a robust, cost-effective approach for the typing of class I HLA alleles in human samples with clinical-grade accuracy.

There are thousands of unique HLA proteins expressed in the human population. These diverse proteins present antigens on the cell surface for immune recognition and constitute the major barrier to allogeneic transplantation. HLA typing is critical for the evaluation of immunological compatibility between organ donor and recipient pairs. Although rapid, high-resolution HLA typing would be ideal for organ allocation^12^, ^13^, this has not been possible due to the technical limitations inherent to both Sanger and second-generation sequencing platforms.

The sequences of all known HLA alleles are deposited in the IPD-IMGT/HLA database^14^. Each HLA allele is named by locus (e.g. *HLA-A*) followed by an asterisk and up to four, colon-delimited numeric fields (Fig. 1a). The first field groups together HLA alleles that encode antigens sharing key serological epitopes. The first and second fields describe groups of alleles that encode the same unique protein. If synonymous mutations are present in any exons, a third field is appended to the allele name, and a fourth field can be added to describe sequence variation in non-coding regions. This comprehensive nomenclature system also allows new alleles to be named and organized hierarchically as they are discovered over time.

**Figure 1.**
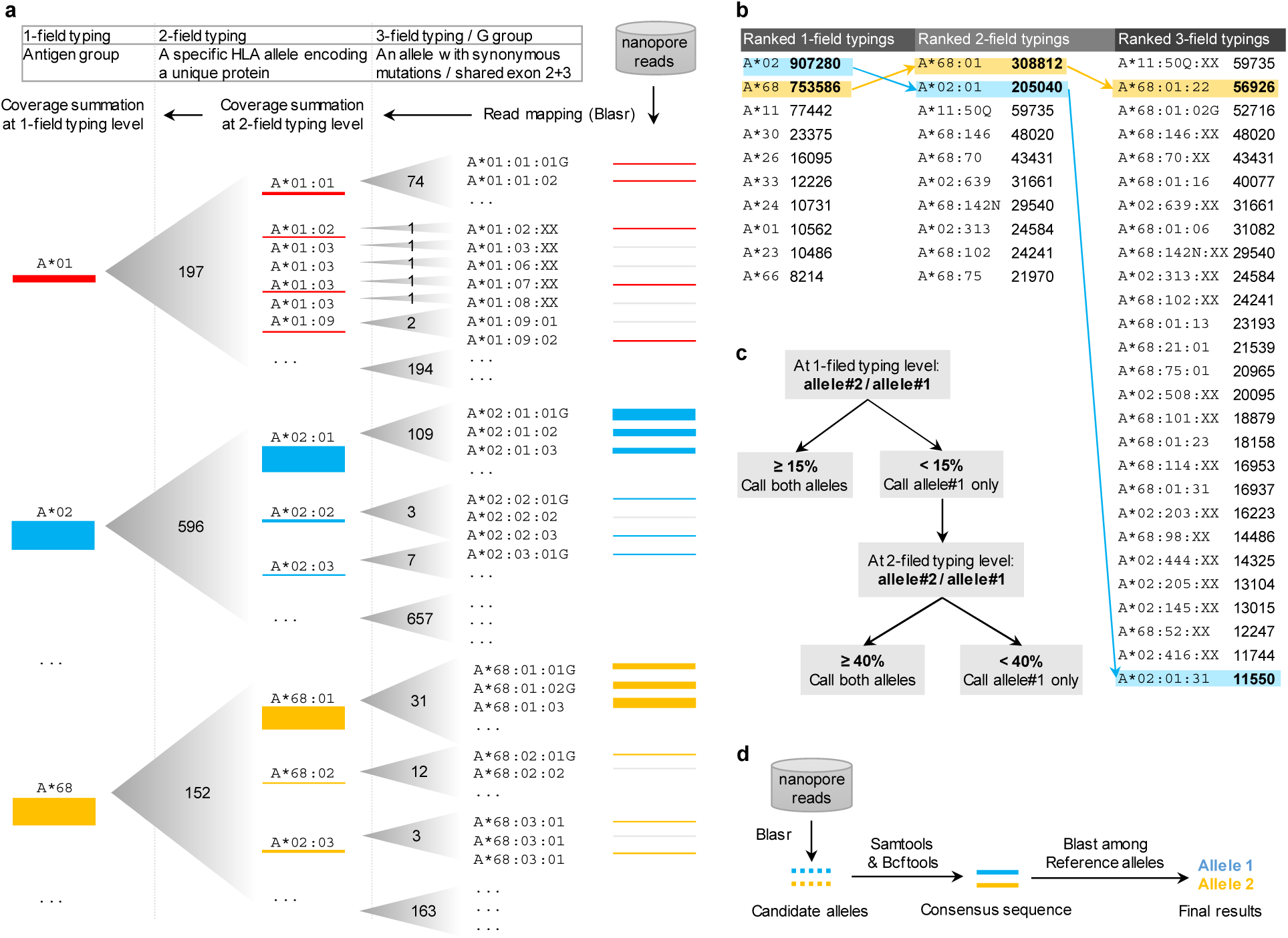
Athlon pipeline for HLA typing by nanopore sequencing. (a) Hierarchical mapping of reads to HLA alleles. Reads are mapped to individual alleles at the 3-field, G-group level (leaves) and coverage is summed to obtain values for each node at the 2- and 1-field level. The hierarchy of the HLA nomenclature system is summarized in the top panel. The numbers in the gradient triangles indicate the total number of 2-field nodes and 3-field leaves. Red, blue and yellow bars represent nanopore reads mapped to different leaves and nodes, and the thickness of the bars represents the depth of coverage. (b) Rank lists of 1-, 2- and 3-field alleles based on the summed total depth of coverage (see Online Methods). Arrows indicate the process of identifying top-ranked, 1-, 2- and 3-field candidate alleles, which are shaded in blue and yellow for a representative heterozgyous sample. (c) An algorithm for calling homozygous versus heterozygous genotypes at the 1- and 2-field typing levels based on the coverage depth of the 2^nd^-ranked allele as a percentage of that of the top-ranked allele. Thresholds of 15% and 40% at the 1-field and 2-field typing levels respectively were established using the homozygous samples from the training dataset. (d) Consensus-based error correction and blasting for final alleles.

The antigen recognition domains (ARD) of class I HLA genes are encoded by exons 2 and 3, the most diverse regions of these genes. These two exons are the most informative for matching donor and recipient, so for hematopoietic stem cell transplantation, HLAs are typed and matched at least at these exons. This is referred to as the G-group level typing.^15^ For solid organ transplantation, HLAs are routinely typed at 1-field resolution, but typing at a higher resolution such as the 3-field, G-group level will provide added benefit.^12^ Therefore, to evaluate the performance of MinION-based HLA typing, we defined success as correctly identifying the one (for homozygous samples) or two (for heterozygous samples) alleles at the 3-field, G-group level that best matched the sequencing data.

We generated three datasets for this study (Online Methods and Supplementary Table 1), by amplifying class I HLA genes using long-range PCR with or without sample barcodes, followed by ligation of sequencing adaptors and multiplexed MinION sequencing. The consensus sequences from the template and complement strands (2D reads) from each run were demultiplexed based on sample barcodes or primer sequences before analysis (Online Methods and Supplementary Fig. 1). Average read lengths consistent with the predicted amplicon sizes ranging from 3.0 to 4.3 kb were achieved across these datasets (Supplementary Table 2). We used one of the datasets that was generated with the earlier R7.3 flow cells (WASHU-T) to develop and train the Athlon pipeline. The two remaining datasets, WASHU-V and DKMS, which were generated with the improved MK1 R7.3 and R9.4 flow cells respectively, were used to evaluate the performance of Athlon.

Since HLA alleles are extremely diverse, mapping reads to any single reference sequence will result in the loss of relevant reads that differ significantly from the chosen reference. To circumvent this problem, Athlon performs two rounds of read mapping to identify candidate alleles and then build consensus sequences from these alleles. First, Athlon maps locus-specific reads to all HLA alleles at the 3-field, G-group resolution (Online Methods) using BLASR, an aligner originally designed to map long and error-prone reads from the PacBio platform.^16^ One or two candidate alleles are then identified based on coverage statistics and an algorithm outlined below (see Fig. 1a-c). Second, all locus-specific reads are realigned to the candidate alleles to generate one or two consensus sequences, which are then queried against all reference alleles in the database to identify the best match as the final typing result (Fig. 1d).

To identify the candidate alleles used to generate the final consensus sequences, Athlon represents each HLA locus by a tree structure, treating the typing fields as branching nodes and leaves in order to identify alleles with the highest read coverage (Fig. 1a). For example, the HLA-A locus has 3311 leaves, 2546 nodes, and 21 nodes at the 3-, 2-, and 1-field typing levels respectively (Fig. 1a). Read coverage at the leaves under each node is summed to provide a total coverage value for each 2-field node and then for each 1-field node. The nodes and leaves are then sorted at each level by total coverage in descending order (Fig. 1b). Next, candidate alleles are identified by first selecting the top-ranked nodes at the 1-field level and then selecting the highest-ranked 2-field nodes that are connected to these 1-field nodes. The highest-ranked 3-field leaves that are connected to the selected 2-field nodes are then chosen as the candidate alleles (Fig. 1b). For the last step of Athlon, all reads are remapped to the selected candidate alleles, and the final consensus sequences are blasted against the reference database to identify the closest allele(s) as the final typing result (Fig. 1d).

To differentiate homozygous versus heterozygous genotypes, Athlon uses cutoffs based on normalized coverage to determine whether to consider a second allele at the one-field and two-field levels. The optimal cutoff values were empirically determined using the WASHU-T training dataset (n=15), which included three homozygous and 27 heterozygous samples. We quantified the coverage of 2^nd^-ranked 1-field and 2-field nodes in the three homozygous samples, and the coverage data were normalized using values from the highest ranked nodes as denominators (Online Methods and Supplementary Fig. 2). The mean plus two standard deviations of the normalized coverage of 2^nd^-ranked nodes was approximately 15% and 40% of the top-ranked allele, which were used as the statistical thresholds for calling a second node (i.e. a heterozygous typing call) at the 1-field and 2-field resolutions respectively (Fig. 1c). With these values for coverage thresholds, we were able to successful classify all homozygous and heterozygous samples in the training dataset. No threshold was applied at the 3-field level because alleles that are heterozygous beyond the first two fields encode the same protein.

Although the training dataset was collected with earlier versions of R7.3 chemistry, which is significantly more error prone than the later R7.3 and current R9.4 chemistry, we achieved typing results that were 96.7%, 83.3%, and 70% concordant with the truth at 1-, 2-, and 3-field resolutions respectively (Fig. 2a). We next used Athlon to type another low accuracy dataset previously published by Ammar et al., which also used the earlier R7.3 chemistry (n=2, a total of 4 alleles typed at 2-field resolution). The Athlon pipeline correctly typed three alleles at the 2-field level (a 75% accuracy), while the original analysis based on the GATK HLACaller was completely discordant at the same resolution (Fig. 2b), demonstrating that Athlon significantly improves HLA typing accuracy over the only existing method.

**Figure 2.**
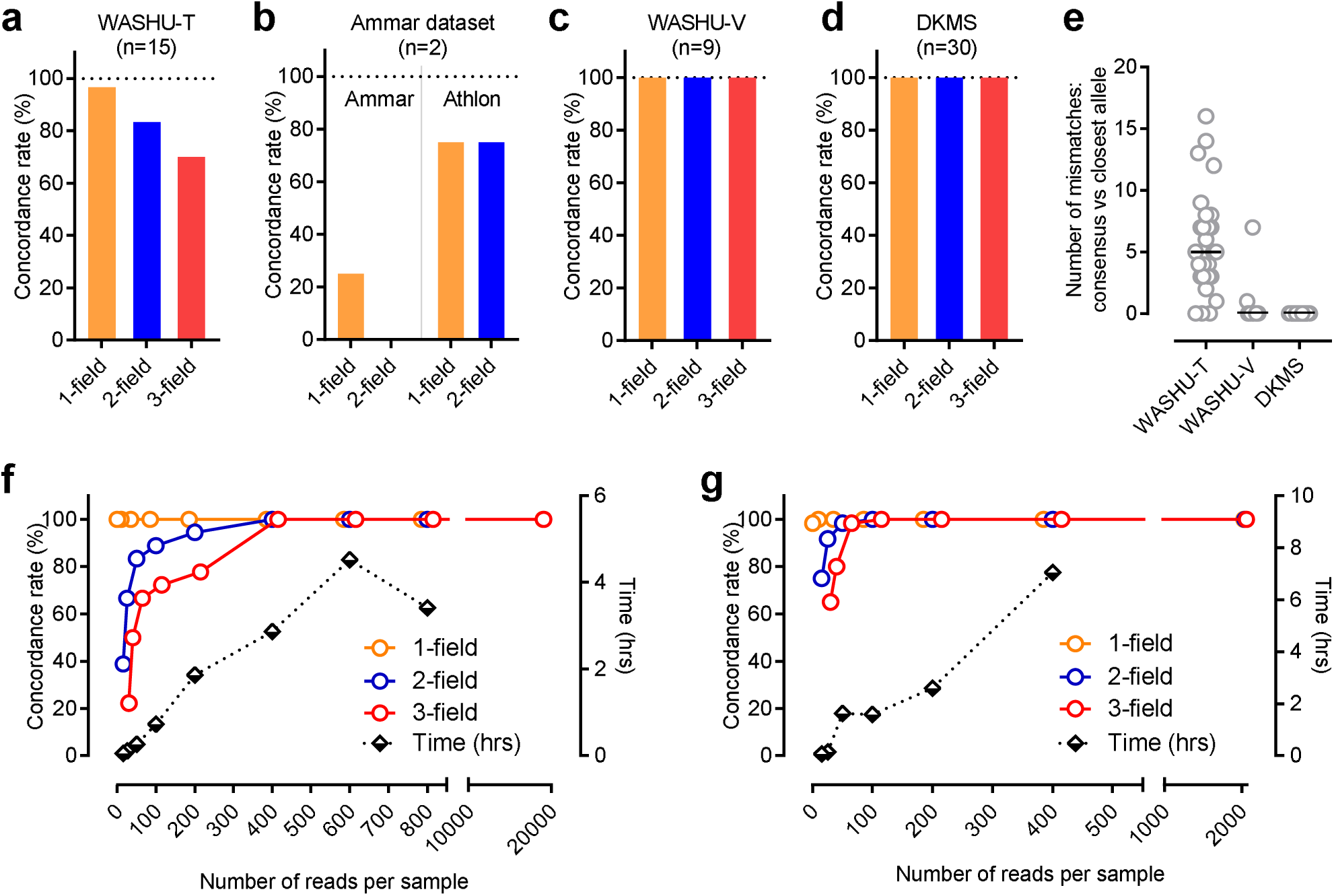
HLA typing accuracy and read downsampling. (a-d) Concordance of HLA typing results for various datasets. (e) Number of mismatches between the predicted consensus sequence and the closest reference allele per sample in the WASHU-T, -V and DKMS datasets. Horizontal bars are medians. (f,g) Impact of down-sampling of the WASHU-V (f) and DKMS (g) datasets on the concordance rates. Concordance at 1-, 2- and 3-field resolutions are plotted against the numbers of input reads for the Athlon pipeline. The computation time is also indicated for different numbers of input reads.

We next sought to evaluate the performance of Athlon using samples sequenced with updated R7.3 and R9.4 flow cells (Supplementary Table 1). We first analyzed the WASHU-V dataset (n=9, including four homozygous samples). For these samples, Athlon was 100% accurate at all three resolutions (Fig. 2c). To validate these results on a larger number of samples, we analyzed the DKMS dataset (n=30), which was collected from a single multiplexed MinION run. Again, the final allele calls were 100% concordant with the ground truth at 1-, 2-and 3-field resolutions (Fig. 2d). Additionally, the number of mismatches between the final predicted consensus sequence(s) and the closest reference allele(s) was quite low for the samples in these two high-quality datasets relative to the earlier, error prone, WASHU-T dataset (Fig. 2e). These results demonstrate that the Athlon pipeline performs HLA typing with clinical-grade accuracy when samples are sequenced with updated R7.3 and R9.4 flow cells.

To determine the maximum number of samples that can be multiplexed on a flow cell, we down-sampled the number of MINION reads used for our analysis and investigated the impact on Athlon’s accuracy. We explored a range of 15-800 reads per sample for the WASHU-V dataset and a range of 15-400 reads per sample for the DKMS dataset. For the WASHU-V dataset, Athlon was 100% accurate at the 1-field level throughout the range of down-sampling. At the 2- and 3-field resolutions, Athlon was 100% concordant if 400 or more reads were input per sample (Fig. 2f). For the DKMS dataset, Athlon was 100% correct at the 1-field level with as few as 25 reads per sample, and only 100 reads per sample were required for a 100% concordance at the 2- and 3-field resolutions (Fig. 2g). These results suggest that up to 600 loci can be typed on a single flow cell, assuming that least 60K total reads can be obtained from each sequencing run (Supplementary Table). Extrapolating from these results, we estimate that more than 100 individuals can be typed for class I HLA genes using a single MinION flow cell. Furthermore, when fewer reads are used per sample, significant reductions in computation time can be achieved (Fig.2f,g, and Online Methods). Taken together, these results suggest that rapid, point-of-care clinical sequencing can be performed cost-effectively through sample multiplexing.

In summary, we have developed and validated a MinION-based method that allows for the accurate typing of highly polymorphic class I HLA genes. One-hundred percent accuracy was obtained for 3-field G-group level typing, a resolution that is suitable for clinical applications. This approach paves the way for point-of-care HLA typing, which would greatly expedite organ allocation since it could be initiated at the bedside of deceased donors, or in an outreach laboratory. Additionally, the library preparation for the MinION does not require the fragmentation of long amplicons, which is regularly applied to HLA typing with second-generation sequencing platforms. Bypassing this step simplifies the workflow, reduces bias,^17^ and results in a more uniform read coverage (Supplementary Fig. 3). Moreover, nanopore sequencing can be readily scaled up to much larger numbers of pores^18^ and does not require a significant capital investment for equipment. In conclusion, MinION-based sequencing has the potential to spark a paradigm shift in HLA typing in contemporary clinical and research laboratories.

## METHODS

Methods, including statements of data availability and any associated references, are available in the online version of the paper.

## ACKNOWLEDGMENTS

We thank Vineeth Surendranath and Vinzenz Lange for their helpful discussion and manuscript suggestions. C. Liu is support by the Washington University Hematology Scholars K12 award (K12-HL087107-07 to C.L.). R. Mitra is supported by grants from the NIH (U01MH109133, R01NS076993) and Children’s Discovery Institute (MC-II-2016-533).

## AUTHOR CONTRIBUTIONS

Contribution: C.L. designed and conducted experiments, constructed the bioinformatic pipeline, analyzed and interpreted the data, and wrote the paper; F.X. performed data analysis and interpretation, worked on an alternative method; J.H. conducted experiments; K.L. conducted experiments and analyzed data; P.Q. analyzed data; B.D. contributed to specimen collection and experiments; R.M. supervised the overall project, designed experiments, analyzed the data, and wrote the paper.

## ONLINE METHODS

### Datasets and study design

Three datasets, WASHU-T (for Washington University-Training), WASHU-V (for validation) and DKMS (for German Bone Marrow Donor Center), were generated and analyzed in this study (Table S1). The WASHU-T dataset used five genomic DNA specimens, four from the 1000 Genomes projects and one from an deidentified homozygous donor, to prepare five multiplexed libraries. Each library included fragmented amplicons from three HLA class I genes (see below) and was sequenced by the R7.3 flow cell chemistry in one run. The WASHU-T dataset was used to develop and train the Athlon pipeline.

The WASHU-V dataset used three genomic DNA specimens, two from the 1000 Genomes projects and one from a deidentified homozygous donor, to generate three libraries. Each multiplexed fragmented amplicons from three HLA class I genes and was sequenced by the updated MK1 R7.3 flow cell chemistry in one run. An external validation was performed using the DKMS dataset, which was generated by multiplexed sequencing of 30 barcoded locus-specific samples in one run using the R9.4 flow cell chemistry. Detailed information including software and protocol versions are provided in Supplementary Table 1. All specimens in the WASHU-T, -V and DKMS datasets were typed by one or more reference methods, including Sanger and Illumina sequencing. Moreover, to benchmark against a method reported previously by Ammar et al.,^19^ the dataset from that report, which included two locus-specific samples (Supplementary Table 1), were analyzed using the Athlon pipeline. The typing results were compared with the truth and typing results reported from the original study. Only 2D reads were analyzed for all datasets due to their lower error rate.

### Target amplification

Three class I HLA genes, HLA-A, -B, and -C, were amplified separately in full length by long-range PCR. Primer sequences are listed in Supplementary Table 1. The primers and PCR conditions used for the WASHU datasets were reported previously.^20^ Long range PCR of the DKMS samples was performed in 96 well plates with 4 µl template DNA, 12.5 µl 2x GoTaq® Long PCR Master Mix (Promega, Madison, USA) and 1 µl of a target-specific primer mix (50 µM each) in a total volume of 25 µl. A thermal profile of 95°C for 3 min followed by 25 cycles at 95°C for 15 s, 62°C for 30 s and 68°C for 7.5 min, and a finishing step at 68°C for 15 min was used. A subsequent PCR was applied for barcoding where a 5 specific adaptor sequence of the target-specific primers was used as template for the barcode introducing primers. This PCR was performed in 96 well plates where 2 µl of the target specific PCR (PCR1) was mixed with 12.5 µl 2x GoTaq® Long PCR Master Mix (Promega, Madison, USA), 8.5 µl of Nuclease-Free Water (Promega, Madison, USA) and 1 µl of an index primer mix (20 µM each); a thermal profile of 95°C for 3 min followed by 7 cycles at 95°C for 15 s, 55°C for 30 s and 68°C for 7.5 min, and a finishing step at 68°C for 15 min was used. Target-specific and indexing primers were designed in-house. All primers were obtained from metabion (metabion international AG, Planegg, Germany).

### Library preparation and nanopore sequencing

For the WASHU-T and -V datasets, PCR amplicons of HLA-A, -B, and -C from the same genomic DNA source were pooled and purified with a 1.0x reaction with Agencourt Ampure XP Beads (Fisher Scientific NC9959336). DNA was then treated with PreCR Repair Mix (NEB M0309) to repair any damaged template DNA. Following a second round of 1x ampure clean-up, libraries were prepared with the standard protocol supplied by ONT using the version MN004 or MN006 library preparation kits. Libraries #1 through #5 were sequenced for 48 hours on the R7.3 flow cells on the original MinION devices (WASHU-T dataset). Libraries #6, #7, and #3 (repeat) were sequenced for 12 hours on the MAP103 (R7.3) flow cells on the newer MK1 MinIONs (WASHU-V dataset). Sequencing and base calling were performed using the MinKnow and Metrichor software available at the time (Supplementary Table 1), and the native FAST5 files were converted to the fastq format using Poretools.^21^

For the DKMS dataset all barcoded amplicons were pooled at 10 µl each and purified initally using SPRIselect Beads (Beckham Coulter, Brea, USA) in a ratio of 0.7:1 beads to PCR product. 1.5 µg of the purified pool was used for ONT library preparation with the ligation based 2D Library Preparation Kit (SQK-LSK208) in combination with an R9.4 SpotON Flowcell (FLO-MIN106). Library Preparation took place according to the manufacturer’s protocol and sequencing was done for 48h on a MK1B MinION. Sequencing and base calling were performed using MinKNOW version 1.1.21 and Metrichor version 1.125. The native Fast5 files were converted to FASTQ files using Poretools.^21^

### Demultiplexing

The WASHU-T, -V and Ammar datasets were demultiplexed based on the locus-specific primer sequences. The python script is provided in the supplementary package online (<u>http://github.com/cliu32/Athlon</u>). The DKMS dataset was demultiplexed using lastlopper.pl (<u>https://github.com/gringer/bioinfscripts/blob/master/lastlopper.pl</u>), a wrapper script for the LAST aligner (<u>http://last.cbrc.jp/</u>)

### HLA nomenclature and result evaluation

HLA alleles are named by the gene name (e.g. HLA-A) followed by an asterisk and up to four digital fields separated by colons. One-field typing corresponds to a group of alleles carrying one or more shared serological antigens. HLA alleles encoding the same unique protein share the same 2-field typing. If synonymous mutations are present in the exons, a third field is appended to the shared 2-field typing to form unique 3-field typings (Figure 1A). For alleles sharing the same 3-field typing but with sequence variations in the non-coding regions, a fourth field is added to form unique 4-field typings. The G-group code was used to describe alleles with identical sequences across exon(s) encoding the peptide binding domains (exons 2 and 3 for class I and exon 2 for class II alleles). To represent a group of such alleles, a capitalized "G" is suffixed to the allele designation of the lowest numbered allele to form the group name, e.g. A*01:01:01G.

For solid organ transplantation, HLAs are largely typed at 1-field resolution as a standard of care, although typing at a higher resolution may have added benefit.^12^ For hematopoietic stem cell transplantation, HLAs are at least typed at the G-group level of resolution.^15^ With these considerations, we aimed to type HLAs at up to the 3-field resolution using the G-group nomenclature. The reference sequences were constructed accordingly to serve this goal (see below). For the WASHU-T, -V, and DKMS datasets, all samples were typed at 3- to 4-field levels by reference methods, the accuracy of nanopore typing results were evaluated by the concordance rates at 1-field, 2-field, and 3-field G-group levels respectively. For the Ammar dataset, because the samples were typed at 2-field level by the reference method, the accuracy of nanopore typing results were evaluated by concordance rate among all alleles at 1- and 2-field levels respectively.

### Reference sequences

Reference sequences for this study were constructed based on the IPD-IMGT/HLA database release 3.26.0.^14^ The .dat file was parsed using Biopython, and sequences for exons 2 and 3 were joint by a large intronic gap filled by "-" as space holders. The final reference file included 3311, 4163, and 3034 sequences for the HLA-A, -B and -C loci, respectively. A separate set of reference sequences without the intronic gap were used for the second read mapping to candidate alleles followed by consensus generation and blast.

### Main procedures and components of the Athlon pipeline

The rationale and workflow of the pipeline are detailed in the Results section. A script for the pipeline, reference files, and sample data are provided at <u>http://github.com/cliu32/Athlon</u>. All read mapping were performed by BLASR (version 2.0.0)^16^ with the “hitPolicy” set as “randombest”. Read coverage was quantified using Bedtools (version 2.25.0) ^22^ with the default setting. Samtools (samtools and bcftools, version 1.2 using htslib 1.2.1)^23^ was used with default setting to manipulate bam files and generate consensus sequences. Final allele call was made using Blast (version 2.2.31)^24^ with the default setting to identify the closest reference allele to a consensus sequence.

### Downsampling experiment

The WASHU-V and DKMS datasets were downsampled to the number of reads per sample as indicated using the seqtk package version 1.0-r31 (Li, <u>https://github.com/lh3/seqtk</u>) followed by analysis using the Athlon pipeline.

### Calculation of quality metrics

The coverage depth at each base position was obtained using Bedtools (Version 2.25.0) for every candidate allele in the DKMS dataset, which was downsampled to 400 reads per sample. The coverage was visualized for each locus. The uniformity of coverage was represented by the coefficient of variance (CV) calculated as the standard deviation of coverage depth at each position across exons 2 and 3 divided by the mean coverage. The allelic balance was calculated as the total coverage of the minor candidate allele divided by that of the major candidate allele in each sample (n=30).

### Data availability

Raw nanopore reads are available from the authors upon request.

## SUPPLEMENTARY MATERIALS

**Supplementary Figure 1.**
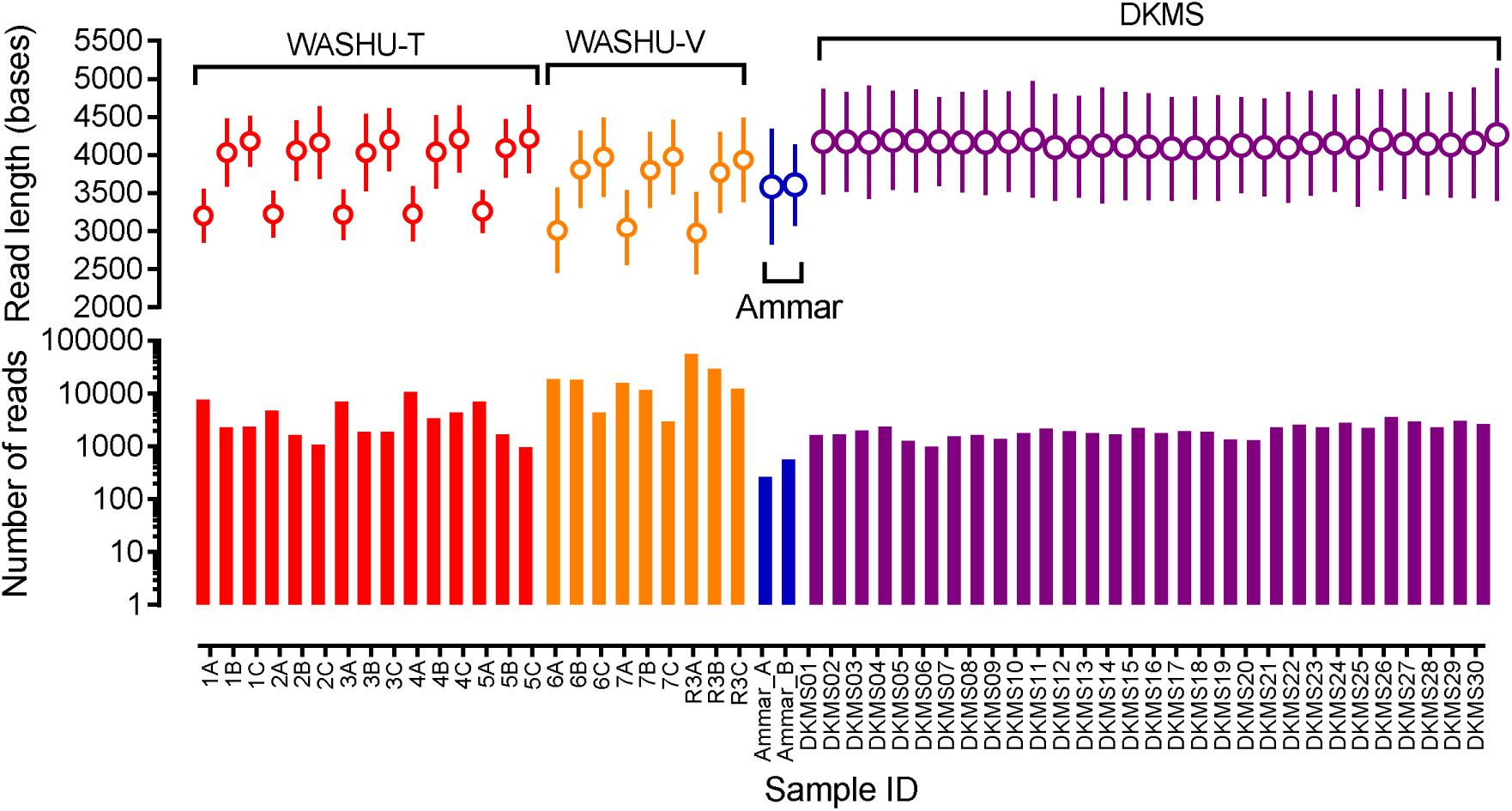
Number of reads and read length in the datasets analyzed for this study. Datasets include WASHU-T (n=15), WASHU-V (n=9), Ammar dataset (n=2), and DKMS dataset (n=30). (Upper panel) Mean read length and standard deviation are shown for each sample. (Lower panel) Mean number of reads are plotted for each sample.

**Supplementary Figure 2.**
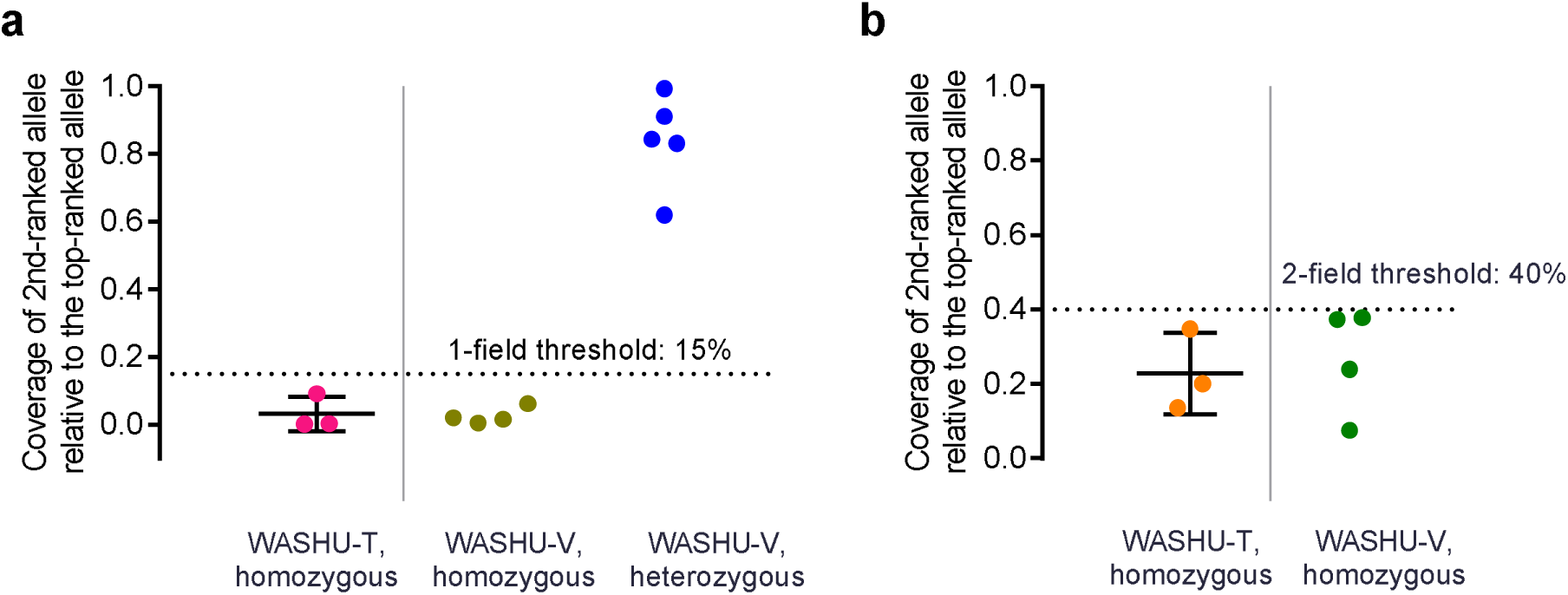
Statistical heterozygosity thresholds at 1-field and 2-field levels to distinguish heterozygous versus homozygous samples. The ratios of the coverage of 2^nd^-ranked allele to that of the top-ranked allele were plotted for the three homozygous samples in the WASHU-T dataset at 1-field (a) and 2-field (b) levels respectively. Horizontal bars are mean (long) and standard deviations (short). Dotted horizontal lines are heterozygosity thresholds, 15% and 40%, which approximated the means plus two standard deviations at 1-field (a) and 2-field (b) levels respectively. The coverage ratios between the 2^nd^- and top-ranked alleles were also plotted for the four homozygous samples and five heterozygous samples in the WASHU-V dataset at 1-field level (a). The ratios for the four homozygous samples at 2-field level were plotted in (b).

**Supplementary Figure 3.**
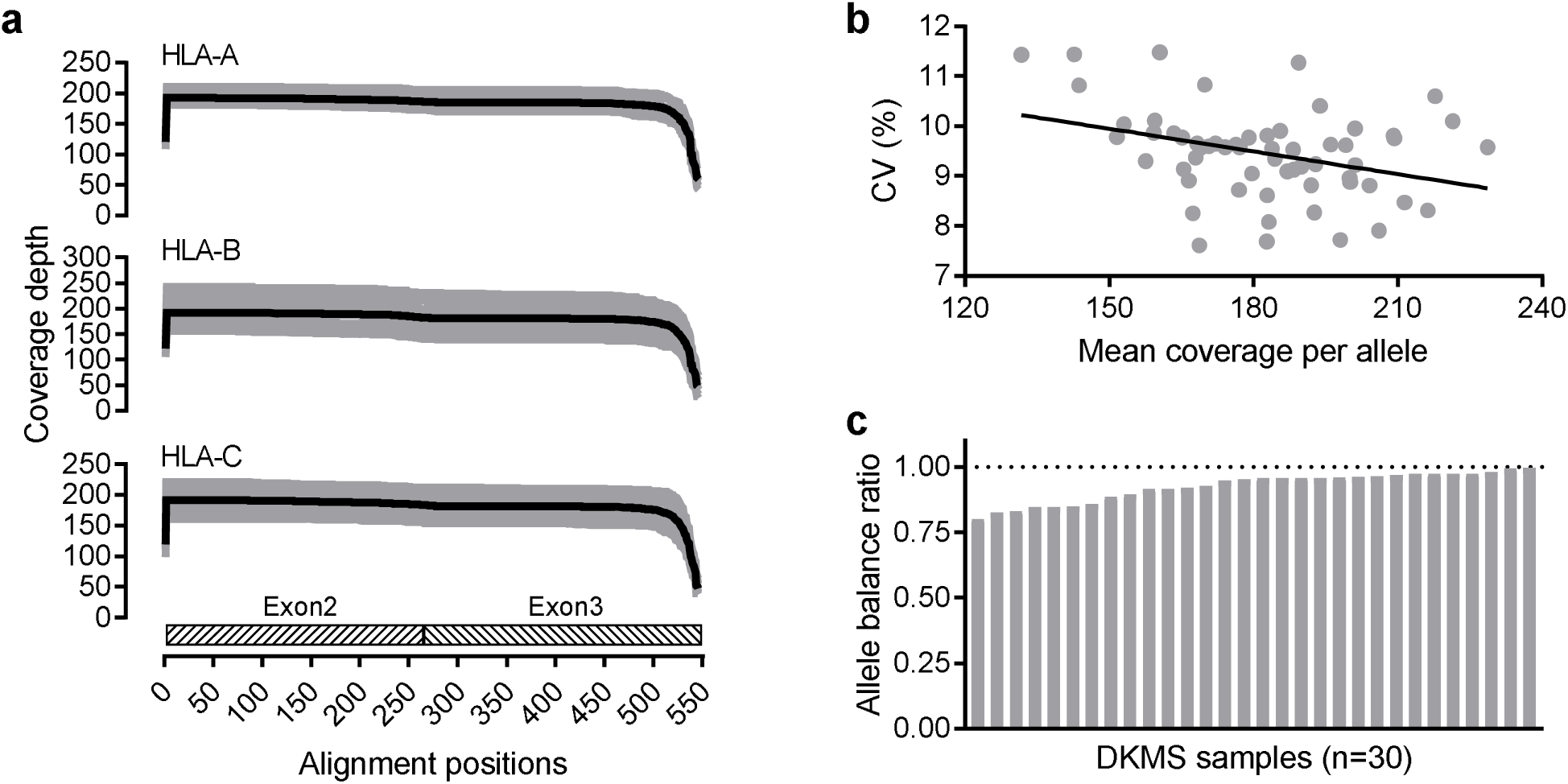
Coverage-related quality metrics. (a) The coverage plots for all candidate alleles from the 30 DKMS samples downsampled to 400 reads per sample are visualized for HLA-A, -B and -C genes (gray lines) in the upper, middle and lower panels respectively. The mean coverage plots (black lines) are superimposed on individual plots. (b) The coefficient of variance (CV) of coverage depths at all positions across a candidate allele, and it correlation with mean coverage depths of individual candidate alleles. A linear regression line is shown. (c) Distribution of allele balance ratios, calculated as the total coverage of the minor allele divided by that of the dominant allele, across 30 DKMS samples.

**Supplementary Table 1.**
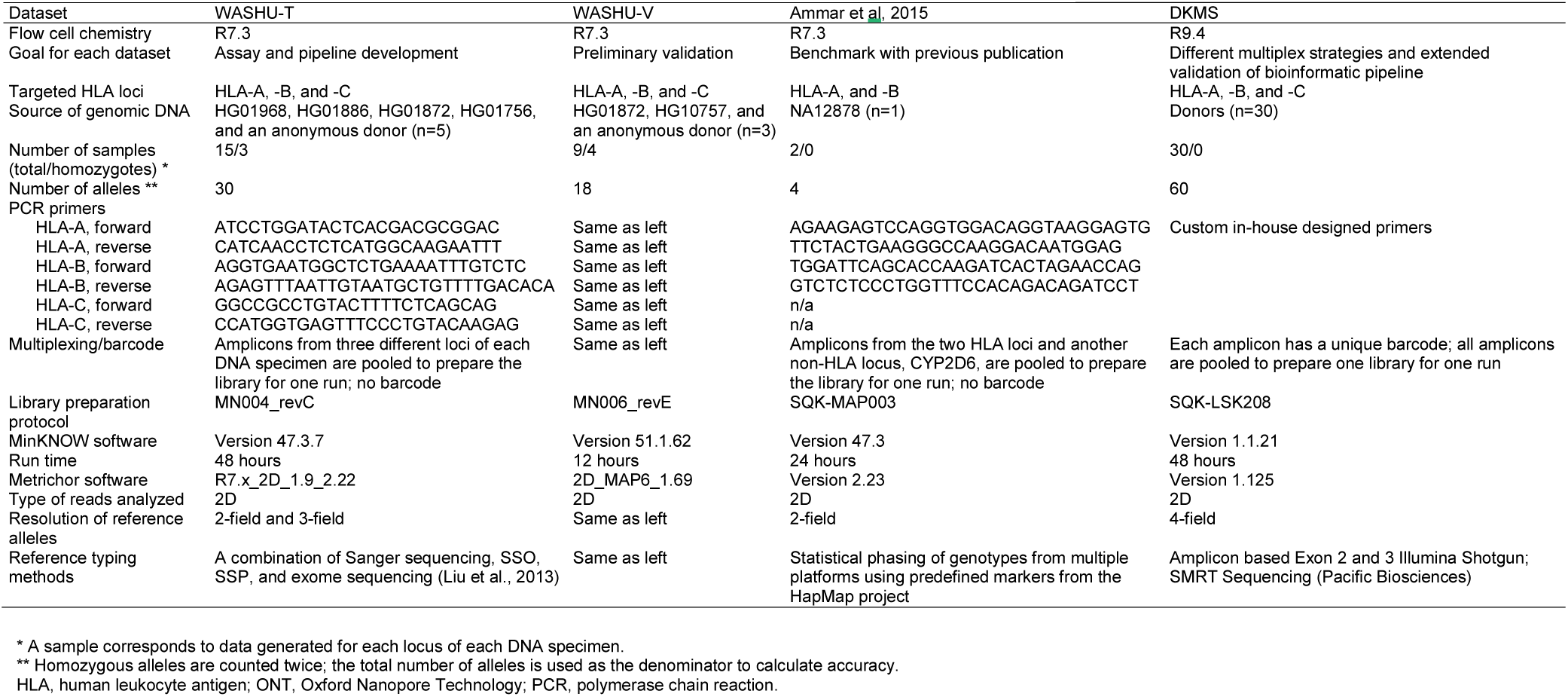
Characteristics of three datasets included in this study.

**Supplementary Table 2.**
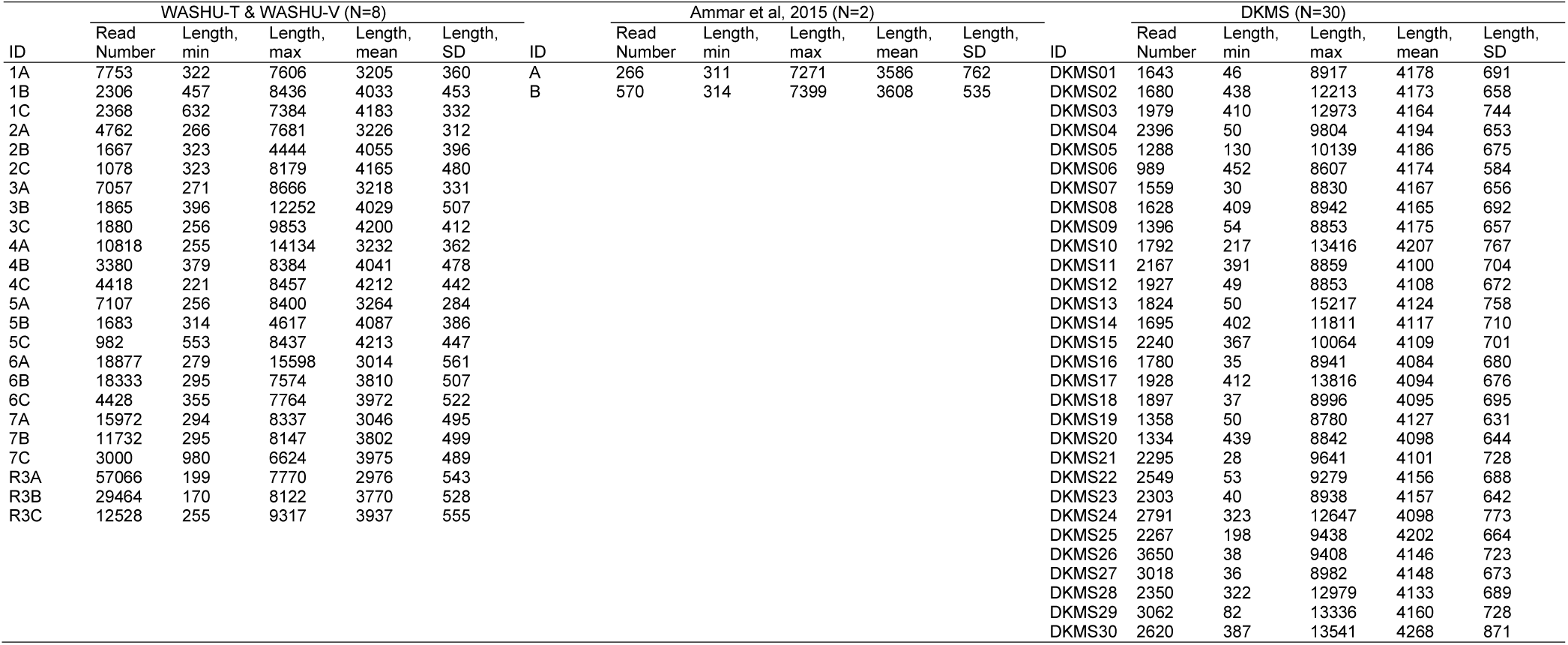
Number and length of 2D reads from the three datasets.

| WASHU-T & WASHU-V (N=8) |  |  |  |  |  | Ammar et al, 2015 (N=2) |  |  |  |  |  | DKMS (N=30) |  |  |  |  |  |
| --- | --- | --- | --- | --- | --- | --- | --- | --- | --- | --- | --- | --- | --- | --- | --- | --- | --- |
| ID | Read Number | Length, min | Length, max | Length, mean | Length, SD | ID | Read Number | Length, min | Length, max | Length, mean | Length, SD | ID | Read Number | Length, min | Length, max | Length, mean | Length, SD |
| 1A | 7753 | 322 | 7606 | 3205 | 360 | A | 266 | 311 | 7271 | 3586 | 762 | DKMS01 | 1643 | 46 | 8917 | 4178 | 691 |
| 1B | 2306 | 457 | 8436 | 4033 | 453 | B | 570 | 314 | 7399 | 3608 | 535 | DKMS02 | 1680 | 438 | 12213 | 4173 | 658 |
| 1C | 2368 | 632 | 7384 | 4183 | 332 |  |  |  |  |  |  | DKMS03 | 1979 | 410 | 12973 | 4164 | 744 |
| 2A | 4762 | 266 | 7681 | 3226 | 312 |  |  |  |  |  |  | DKMS04 | 2396 | 50 | 9804 | 4194 | 653 |
| 2B | 1667 | 323 | 4444 | 4055 | 396 |  |  |  |  |  |  | DKMS05 | 1288 | 130 | 10139 | 4186 | 675 |
| 2C | 1078 | 323 | 8179 | 4165 | 480 |  |  |  |  |  |  | DKMS06 | 989 | 452 | 8607 | 4174 | 584 |
| 3A | 7057 | 271 | 8666 | 3218 | 331 |  |  |  |  |  |  | DKMS07 | 1559 | 30 | 8830 | 4167 | 656 |
| 3B | 1865 | 396 | 12252 | 4029 | 507 |  |  |  |  |  |  | DKMS08 | 1628 | 409 | 8942 | 4165 | 692 |
| 3C | 1880 | 256 | 9853 | 4200 | 412 |  |  |  |  |  |  | DKMS09 | 1396 | 54 | 8853 | 4175 | 657 |
| 4A | 10818 | 255 | 14134 | 3232 | 362 |  |  |  |  |  |  | DKMS10 | 1792 | 217 | 13416 | 4207 | 767 |
| 4B | 3380 | 379 | 8384 | 4041 | 478 |  |  |  |  |  |  | DKMS11 | 2167 | 391 | 8859 | 4100 | 704 |
| 4C | 4418 | 221 | 8457 | 4212 | 442 |  |  |  |  |  |  | DKMS12 | 1927 | 49 | 8853 | 4108 | 672 |
| 5A | 7107 | 256 | 8400 | 3264 | 284 |  |  |  |  |  |  | DKMS13 | 1824 | 50 | 15217 | 4124 | 758 |
| 5B | 1683 | 314 | 4617 | 4087 | 386 |  |  |  |  |  |  | DKMS14 | 1695 | 402 | 11811 | 4117 | 710 |
| 5C | 982 | 553 | 8437 | 4213 | 447 |  |  |  |  |  |  | DKMS15 | 2240 | 367 | 10064 | 4109 | 701 |
| 6A | 18877 | 279 | 15598 | 3014 | 561 |  |  |  |  |  |  | DKMS16 | 1780 | 35 | 8941 | 4084 | 680 |
| 6B | 18333 | 295 | 7574 | 3810 | 507 |  |  |  |  |  |  | DKMS17 | 1928 | 412 | 13816 | 4094 | 676 |
| 6C | 4428 | 355 | 7764 | 3972 | 522 |  |  |  |  |  |  | DKMS18 | 1897 | 37 | 8996 | 4095 | 695 |
| 7A | 15972 | 294 | 8337 | 3046 | 495 |  |  |  |  |  |  | DKMS19 | 1358 | 50 | 8780 | 4127 | 631 |
| 7B | 11732 | 295 | 8147 | 3802 | 499 |  |  |  |  |  |  | DKMS20 | 1334 | 439 | 8842 | 4098 | 644 |
| 7C | 3000 | 980 | 6624 | 3975 | 489 |  |  |  |  |  |  | DKMS21 | 2295 | 28 | 9641 | 4101 | 728 |
| R3A | 57066 | 199 | 7770 | 2976 | 543 |  |  |  |  |  |  | DKMS22 | 2549 | 53 | 9279 | 4156 | 688 |
| R3B | 29464 | 170 | 8122 | 3770 | 528 |  |  |  |  |  |  | DKMS23 | 2303 | 40 | 8938 | 4157 | 642 |
| R3C | 12528 | 255 | 9317 | 3937 | 555 |  |  |  |  |  |  | DKMS24 | 2791 | 323 | 12647 | 4098 | 773 |
|  |  |  |  |  |  |  |  |  |  |  |  | DKMS25 | 2267 | 198 | 9438 | 4202 | 664 |
|  |  |  |  |  |  |  |  |  |  |  |  | DKMS26 | 3650 | 38 | 9408 | 4146 | 723 |
|  |  |  |  |  |  |  |  |  |  |  |  | DKMS27 | 3018 | 36 | 8982 | 4148 | 673 |
|  |  |  |  |  |  |  |  |  |  |  |  | DKMS28 | 2350 | 322 | 12979 | 4133 | 689 |
|  |  |  |  |  |  |  |  |  |  |  |  | DKMS29 | 3062 | 82 | 13336 | 4160 | 728 |
|  |  |  |  |  |  |  |  |  |  |  |  | DKMS30 | 2620 | 387 | 13541 | 4268 | 871 |

**Supplementary Table 3.** HLA typing results and candidate allele pairs for the WASHU-T and WASHU-V datasets.

| ID | Allele | Dataset | Truth | Final results |  |  |  | Candidate alleles |  |  |  |
| --- | --- | --- | --- | --- | --- | --- | --- | --- | --- | --- | --- |
|  |  |  |  | All reads | 400 reads | 200 reads | 100 reads | All reads | 400 reads | 200 reads | 100 reads |
| 1A | Allele1 | WASHU-T | A*02:01:01 | A*02:119:XX | A*02:01:25 | A*02:01:119 | A*02:01:94 | A*02:01:31 | A*02:01:25 | A*02:01:119 | A*02:01:94 |
| 1A | Allele2 | WASHU-T | A*68:01:02 | A*68:01:22 | A*68:01:22 | A*68:01:22 | A*68:01:22 | A*68:01:22 | A*68:01:22 | A*68:01:22 | A*68:01:22 |
| 1B | Allele1 | WASHU-T | B*07:02:01 | B*07:02:03 | B*07:02:04 | B*07:67N:XX | B*07:02:04 | B*07:02:03 | B*07:02:40 | B*07:67N:XX | B*07:02:04 |
| 1B | Allele2 | WASHU-T | B*40:02:01 | B*40:02:01G | B*40:02:08 | B*40:02:07 | B*40:02:01G | B*40:02:06 | B*40:02:08 | B*40:02:07 | B*40:02:01G |
| 1C | Allele1 | WASHU-T | C*07:02:01 | C*07:02:12 | C*07:02:32 | C*07:02:32 | C*07:02:57 | C*07:02:12 | C*07:02:32 | C*07:02:32 | C*07:02:57 |
| 1C | Allele2 | WASHU-T | C*03:04:01 | C*03:04:01G | C*03:04:38 | C*03:04:38 | C*03:03:05 | C*03:04:16 | C*03:04:38 | C*03:04:38 | C*03:03:05 |
| 2A | Allele1 | WASHU-T | A*30:02:01 | A*30:02:01G | A*30:02:01G | A*30:02:01G | A*30:02:01G | A*30:02:01G | A*30:02:09 | A*30:02:01G | A*30:02:09 |
| 2A | Allele2 | WASHU-T | A*74:01:01G | A*74:01:01G | A*74:01:01G | A*74:01:04 | A*74:01:01G | A*74:01:01G | A*74:01:01G | A*74:01:04 | A*74:01:01G |
| 2B | Allele1 | WASHU-T | B*57:03:01 | B*57:03:01G | B*57:03:01G | B*57:01:01G | B*57:01:08 | B*57:01:01G | B*57:01:01G | B*57:01:03 | B*57:01:08 |
| 2B | Allele2 | WASHU-T | B*15:03:01 | B*15:03:01G | B*15:266:XX | B*15:266:XX | B*15:266:XX | B*15:266:XX | B*15:266:XX | B*15:266:XX | B*15:266:XX |
| 2C | Allele1 | WASHU-T | C*02:10:01G | C*02:02:06 | C*02:02:15 | C*02:02:09 | C*02:02:06 | C*02:02:06 | C*02:02:15 | C*02:02:09 | C*02:02:06 |
| 2C | Allele2 | WASHU-T | C*07:01:02 | C*07:01:01G | C*07:01:17 | C*07:01:17 | C*07:57:XX | C*07:01:17 | C*07:01:17 | C*07:01:17 | C*07:57:XX |
| 3A | Allele1 | WASHU-T | A*24:07:01 | A*24:02:64 | A*24:02:34G | A*24:02:72 | A*24:02:64 | A*24:02:64 | A*24:02:34G | A*24:02:72 | A*24:02:64 |
| 3A | Allele2 | WASHU-T | A*11:02:01 | A*11:02:01G | A*11:02:01G | A*11:02:01G | A*11:01:33 | A*11:02:01G | A*11:02:01G | A*11:02:01G | A*11:01:33 |
| 3B | Allele1 | WASHU-T | B*27:04:01 | B*27:04:01G | B*27:04:01G | B*27:04:01G | B*27:04:01G | B*27:04:01G | B*27:04:01G | B*27:04:01G | B*27:04:01G |
| 3B | Allele2 | WASHU-T | B*39:05:01 | B*39:05:01 | B*39:01:18 | B*39:01:18 | B*39:121:XX | B*39:01:04 | B*39:01:18 | B*39:01:18 | B*39:121:XX |
| 3C | Allele1 | WASHU-T | C*08:01:01 | C*08:01:01G | C*08:01:01G | C*08:01:12 | C*08:01:01G | C*08:01:01G | C*08:01:01G | C*08:01:12 | C*08:01:01G |
| 3C | Allele2 | WASHU-T | C*12:02:02 | C*12:02:01G | C*12:02:01G | C*12:02:09 | C*12:02:09 | C*12:02:01G | C*12:02:01G | C*12:02:09 | C*12:02:09 |
| 4A | Allele1 | WASHU-T | A*30:02:01 | A*30:02:01G | A*30:02:01G | A*30:02:01G | A*30:02:09 | A*30:02:01G | A*30:02:09 | A*30:02:09 | A*30:02:09 |
| 4A | Allele2 | WASHU-T | A*66:01:01G | A*26:01:23 | A*26:01:06 | A*26:01:02 | A*26:17:XX | A*26:01:06 | A*26:01:06 | A*26:01:02 | A*26:17:XX |
| 4B | Allele1 | WASHU-T | B*41:02:01 | B*41:02:01G | B*41:02:01G | B*41:02:01G | B*41:39:XX | B*41:02:01G | B*41:02:01G | B*41:02:01G | B*41:39:XX |
| 4B | Allele2 | WASHU-T | B*18:01:01 | B*18:114:XX | B*18:01:01G | B*18:01:01G | B*18:01:02 | B*18:01:01G | B*18:01:01G | B*18:01:01G | B*18:01:02 |
| 4C | Allele1 | WASHU-T | C*17:01:01 | C*17:01:07 | C*17:01:07 | C*17:01:07 | C*17:01:07 | C*17:01:07 | C*17:01:07 | C*17:01:07 | C*17:01:07 |
| 4C | Allele2 | WASHU-T | C*05:01:01 | C*05:01:01G | C*05:01:01G | C*05:01:02 | C*05:31:XX | C*05:01:01G | C*05:01:02 | C*05:01:02 | C*05:31:XX |
| 5A | Allele1 | WASHU-T | A*01:01:01 | A*01:01:01G | A*01:01:01G | A*01:01:01G | A*01:01:01G | A*01:01:01G | A*01:01:20 | A*01:01:20 | A*01:01:01G |
| 5A | Allele2 | WASHU-T | A*01:01:01 | A*01:01:01G | A*01:01:01G | A*01:01:01G | A*01:01:01G | A*01:01:01G | A*01:01:20 | A*01:01:20 | A*01:01:01G |
| 5B | Allele1 | WASHU-T | B*08:01:01/20 | B*08:01:01G | B*08:01:01G | B*08:01:01G | B*08:01:06 | B*08:01:06 | B*08:01:01G | B*08:01:06 | B*08:01:06 |
| 5B | Allele2 | WASHU-T | B*08:01:01/20 | B*08:01:01G | B*08:01:01G | B*08:01:01G | B*08:01:06 | B*08:01:06 | B*08:01:01G | B*08:01:06 | B*08:01:06 |
| 5C | Allele1 | WASHU-T | C*07:01:01/01 | C*07:01:01G | C*07:01:01G | C*07:01:01G | C*07:01:01G | C*07:01:01G | C*07:01:01G | C*07:01:01G | C*07:01:01G |
| 5C | Allele2 | WASHU-T | C*07:01:01/01 | C*07:01:01G | C*07:01:01G | C*07:01:01G | C*07:01:01G | C*07:01:01G | C*07:01:01G | C*07:01:01G | C*07:01:01G |
| 6A | Allele1 | WASHU-V | A*03:01:01:04 | A*03:01:01G | A*03:01:01G | A*03:01:01G | A*03:01:01G | A*03:01:23 | A*03:01:23 | A*03:01:23 | A*03:01:23 |
| 6A | Allele2 | WASHU-V | A*03:01:01:06 | A*03:01:01G | A*03:01:01G | A*03:01:01G | A*03:01:01G | A*03:01:23 | A*03:01:23 | A*03:01:23 | A*03:01:23 |
| 6B | Allele1 | WASHU-V | B*57:01:01 | B*57:01:01G | B*57:01:01G | B*57:01:01G | B*57:01:01G | B*57:01:01G | B*57:01:01G | B*57:01:01G | B*57:01:01G |
| 6B | Allele2 | WASHU-V | B*57:01:01 | B*57:01:01G | B*57:01:01G | B*57:01:01G | B*57:01:01G | B*57:01:01G | B*57:01:01G | B*57:01:01G | B*57:01:01G |
| 6C | Allele1 | WASHU-V | C*06:02:01:01 | C*06:02:01G | C*06:02:01G | C*06:02:01G | C*06:02:01G | C*06:02:01G | C*06:02:01G | C*06:02:01G | C*06:02:01G |
| 6C | Allele2 | WASHU-V | C*06:02:01:01 | C*06:02:01G | C*06:02:01G | C*06:02:01G | C*06:02:01G | C*06:02:01G | C*06:02:01G | C*06:02:01G | C*06:02:01G |
| 7A | Allele1 | WASHU-V | A*02:01:01 | A*02:01:01G | A*02:01:01G | A*02:01:87 | A*02:01:81 | A*02:01:87 | A*02:01:47 | A*02:01:87 | A*02:01:81 |
| 7A | Allele2 | WASHU-V | A*01:01:01:01 | A*01:01:01G | A*01:01:01G | A*01:01:63 | A*01:01:13 | A*01:01:34 | A*01:01:34 | A*01:01:63 | A*01:01:13 |
| 7B | Allele1 | WASHU-V | B*57:01:01 | B*57:01:01G | B*57:01:01G | B*57:01:01G | B*57:01:09 | B*57:01:01G | B*57:01:09 | B*57:01:01G | B*57:01:09 |
| 7B | Allele2 | WASHU-V | B*18:01:01 | B*18:01:01G | B*18:01:01G | B*18:01:01G | B*18:01:01G | B*18:01:01G | B*18:01:01G | B*18:01:01G | B*18:01:01G |
| 7C | Allele1 | WASHU-V | C*07:01:01:01 | C*07:01:01G | C*07:01:01G | C*07:01:34 | C*07:01:01G | C*07:01:01G | C*07:01:07 | C*07:01:34 | C*07:01:01G |
| 7C | Allele2 | WASHU-V | C*07:01:01:01 | C*07:01:01G | C*07:01:01G | C*07:528:XX | C*07:528:XX | C*07:01:01G | C*07:01:07 | C*07:528:XX | C*07:528:XX |
| R3A | Allele1 | WASHU-V | A*11:02:01 | A*11:02:01G | A*11:02:01G | A*11:02:01G | A*11:02:01G | A*11:02:01G | A*11:02:01G | A*11:02:01G | A*11:02:01G |
| R3A | Allele2 | WASHU-V | A*24:07:01 | A*24:07:01 | A*24:07:01 | A*24:07:01 | A*24:07:01 | A*24:02:41 | A*24:07:01 | A*24:02:17 | A*24:07:01 |
| R3B | Allele1 | WASHU-V | B*27:04:01 | B*27:04:01G | B*27:04:01G | B*27:04:01G | B*27:04:01G | B*27:04:01G | B*27:04:01G | B*27:04:01G | B*27:04:01G |
| R3B | Allele2 | WASHU-V | B*39:05:01 | B*39:05:01 | B*39:05:01 | B*39:05:01 | B*39:01:01G | B*39:01:13 | B*39:01:13 | B*39:31:XX | B*39:01:13 |
| R3C | Allele1 | WASHU-V | C*08:01:01 | C*08:01:01G | C*08:01:01G | C*08:01:01G | C*08:01:06 | C*08:01:06 | C*08:01:06 | C*08:01:06 | C*08:01:06 |
| R3C | Allele2 | WASHU-V | C*12:02:02 | C*12:02:01G | C*12:02:01G | C*12:02:01G | C*12:02:01G | C*12:02:01G | C*12:02:01G | C*12:02:01G | C*12:02:01G |

**Discordance in 1-field typing** | **Discordance in 2-field typing** | **Discordance in 3-field typing** | **Truth and typing result in the same G group**

**Supplementary Table 4.** HLA typing results and candidate allele pairs for the Ammar dataset.

| ID | Allele | Truth | Ammar et al. | Athlon |  |
| --- | --- | --- | --- | --- | --- |
|  |  |  |  | Final results | Candidate alleles |
| Ammar_A | Allele1 | A*01:01 | A*01:32 | A*01:01:73 | A*01:01:73 |
| Ammar_A | Allele2 | A*11:01 | A*03:12 | A*11:01:08 | A*11:01:60 |
| Ammar_B | Allele1 | B*08:01 | B*07:65 | B*08:01:06 | B*08:01:04 |
| Ammar_B | Allele2 | B*56:01 | B*55:10 | B*55:01:01G | B*55:48:XX |
| Discordance in 1-field typing \| Discordance in 2-field typing |  |  |  |  |  |

**Supplementary Table 5.**
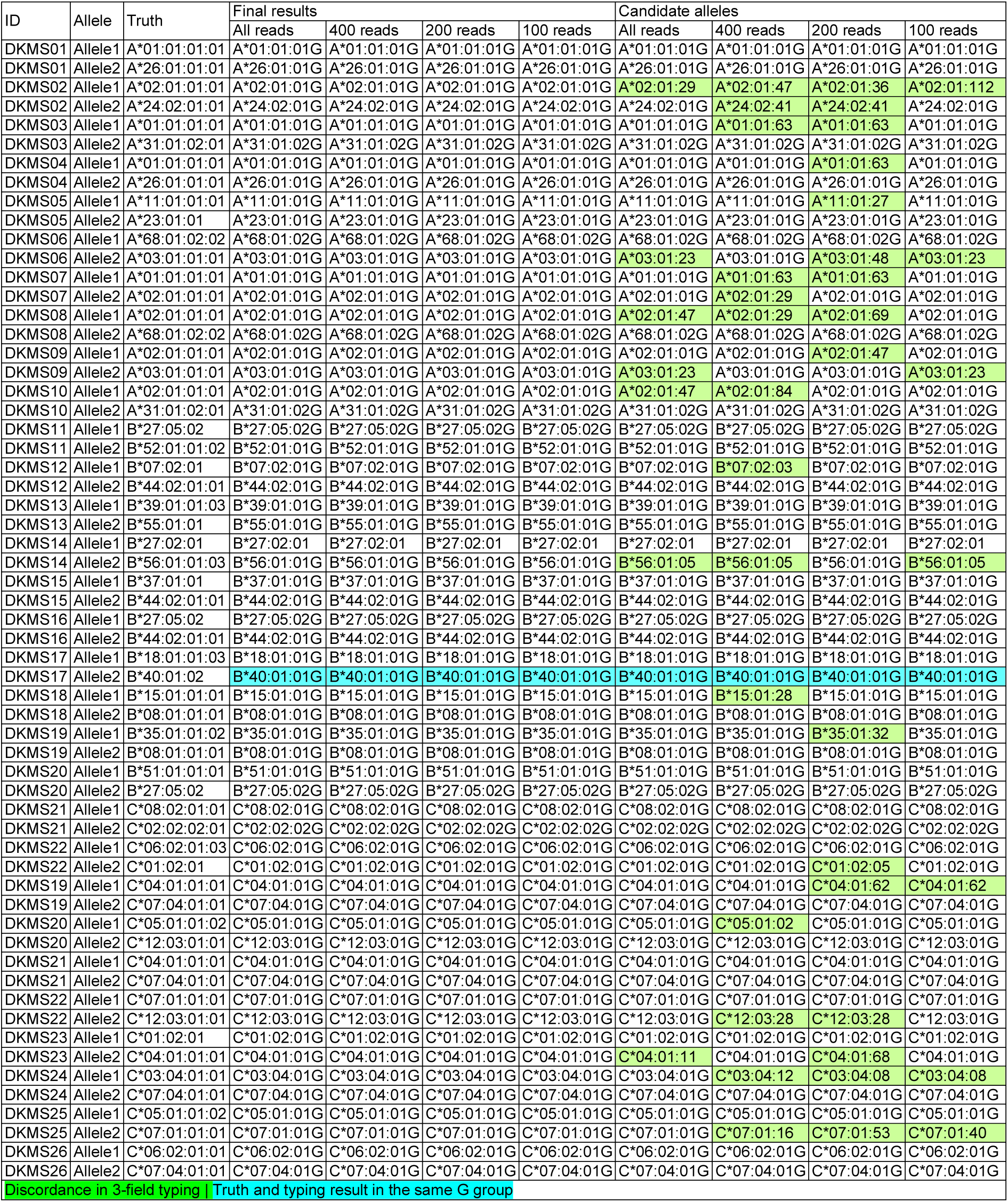
HLA typing results and candidate allele pairs for the DKMS dataset.

